# Novel Spatial-Structural-Zero-Aware Dissimilarity Measures for Subtype Discovery Using Single Cell Hi-C Data

**DOI:** 10.1101/2025.07.23.666407

**Authors:** Yongqi Liu, Victor Jin, Shili Lin

**Author notes:** Address for correspondence: Shili Lin, PhD, Department of Statistics, The Ohio State University, 1958 Neil Avenue, Columbus, OH 43210-1247, USA.

## Abstract

High-throughput single-cell Hi-C (scHi-C) technologies have opened new avenues for investigating cell-to-cell variability in the three-dimensional organization of the genome within individual nuclei. Despite their potential, analyses of scHi-C data are hindered by data sparsity, which varies substantially across cells. To address this challenge, recent methods aim to denoise scHi-C data and differentiate between two types of zero entries: structural zeros (SZs), which reflect true absence of contacts due to biological structure, and dropouts (DOs), which arise from insufficient sequencing depth. However, current dissimilarity measures used in downstream analyses, such as Euclidean distance and Kendall’s tau, treat all zeros as equivalent, thus do not distinguish between SZs and DOs nor recognize the special role of SZs in capturing cell-to-cell variability. Such oversight limits the ability to accurately capture biologically meaningful differences between cells. In this study, we introduce structural-zeros-aware Kendall’s tau (szKendall), a novel dissimilarity metric that explicitly incorporates the spatial structure of 2D contact matrices and leverages the presence or absence of shared SZs across cells. Through comprehensive simulations and analysis of real scHi-C datasets, we demonstrate that szKendall more effectively captures key structural features and achieves superior performance in cell clustering tasks compared to existing approaches. Our results underscore the importance of SZ-aware dissimilarity measures in advancing single-cell 3D genomics.

## 1 Introduction

Over the past 15 years, advances in genomic profiling and computational methodologies have significantly transformed our understanding of three-dimensional (3D) chromatin architecture [1, 2, 3, 4, 5]. The human genome is now recognized as a highly organized structure composed of hierarchical chromatin units, including chromosome territories, compartments, topologically associating domains, and chromatin loops, which contribute to spatial organization and regulatory functions of the genome [6, 7, 8].

The advent of high-throughput single-cell Hi-C (scHi-C) assays since 2017 [9, 10, 11, 12] has raised great expectations for dissecting cell-to-cell variability in genome architecture. However, realizing this promise requires tackling the unique analytical challenges posed by scHi-C data, most notably the extreme sparsity and intrinsic biological variability.

First and foremost, scHi-C data are exceptionally sparse. While data sparsity is a common issue in other single-cell modalities such as single-cell RNA sequencing (scRNA-seq), the problem is even more acute in scHi-C. State-of-the-art single-cell DNA-based assays typically achieve 5 – 10% genome coverage, which translates to only about 1% coverage in the 2D contact matrices used in scHi-C [13]. Second, there is substantial spatiotemporal variability in chromatin structure across individual cells even among cells of the same type, as highlighted by image-based techniques [14, 15]. A contact observed in one cell may be absent in another, exhibiting true biological heterogeneity rather than technical noise.

Together, these challenges complicate the interpretation of zero entries in scHi-C contact matrices. A zero may correspond to a structural zero (SZ), indicating a genuine lack of contact between two loci, or a dropout (DO), arising from insufficient sequencing depth. Disentangling these two sources of zeros is crucial for accurately characterizing cell-to-cell variability and discovering cellular subpopulations.

Addressing these issues, several methods have been developed to improve scHi-C data quality and differentiate between SZs and DOs. Examples include HiCRep [16], which uses a 2D mean filter (MF); SCL [17], which applies a 2D Gaussian kernel (GK); and GenomeDisco [18], based on random walk smoothing (RW). Additional efforts include scHiCluster [13], which integrates random walk with restart and linear convolution, and Higashi [19], which uses a hypergraph-based neural embedding. Model-based approaches have also emerged, such as a latent Dirichlet allocation method for identifying chromatin compartment patterns [20]. More recently, specialized methods like HiCImpute [21] and scHiCSRS [22] explicitly aim to separate structural zeros from dropouts.

Despite these advances, a key limitation remains: downstream analyses such as clustering and subtype discovery typically ignore structural zero information. Most existing dissimilarity measures, such as Euclidean distance and Kendall’s tau, treat all zeros uniformly, overlooking spatial SZ patterns that may carry biologically meaningful signals. We argue that this oversight limits the ability to detect fine-grained structural differences among cells. For example, in the human prefrontal cortex, L4 and L5 excitatory neuron subtypes reside in distinct cortical layers and are expected to be well separated. However, clustering based on Hi-C data from single-nucleus methyl-3C sequencing (sn-m3C-seq) [23] fails to distinguish these subtypes effectively [24]. Heatmaps of raw and smoothed contact matrices (using MF, GK, and RW methods) reveal substantial similarity between subsets of L4 and L5 cells, leading to their misclassification by standard clustering approaches like K-means using Euclidean distance (Supplementary Fig. S1). Such clustering algorithms do not account for SZ information nor spatial structure of the 2D contact matrices.

To tackle this crucial shortfall, we propose a new dissimilarity measure, structural-zeros-aware Kendall’s tau (szKendall), for comparing scHi-C contact matrices across cells. szKendall explicitly incorporates spatial structure and the concordance or discordance of SZ positions between cells, providing a biologically informed way to quantify differences. We conduct extensive simulations to benchmark its performance and apply the method to two real datasets: mouse embryonic stem cells (mESC) [25] and human prefrontal cortex sn-m3C-seq data [23]. Our results demonstrate that szKendall enhances clustering accuracy and reveals subtle yet meaningful chromatin structural differences that are missed by existing methods.

## 2 Results

### 2.1 Overview of szKendall

We introduce a novel dissimilarity measure, spatial-structural-zero-aware Kendall’s tau (szKendall), which modifies the standard Kendall’s tau distance by incorporating information on structural zeros (SZs) and the spatial organization of locus pairs (LPs) in single-cell Hi-C contact matrices.

Standard Kendall’s tau operates on two vectors of equal length. In our setting, these vectors are the vectorized upper triangular portions of scHi-C contact matrices of two cells (SC1 and SC2), representing all LPs (Fig. 1a). For each pair of LPs (LP_*ij*_ and LP_*uv*_), the method evaluates concordance: if SC1 and SC2 agree on the relative ordering of contact values (i.e., the contact count at LP_*ij*_ is larger (or smaller) than that at LP_*uv*_ for both cells), a score of 0 is assigned. On the other hand, discordance — defined opposite of concordance — receives a score of 1; and ties receive 0.5. The average of these scores over all LP pairs yields a Kendall’s tau distance between SC1 and SC2, which ranges from 0 to 1, with larger values indicating greater dissimilarity.

**Fig. 1:**
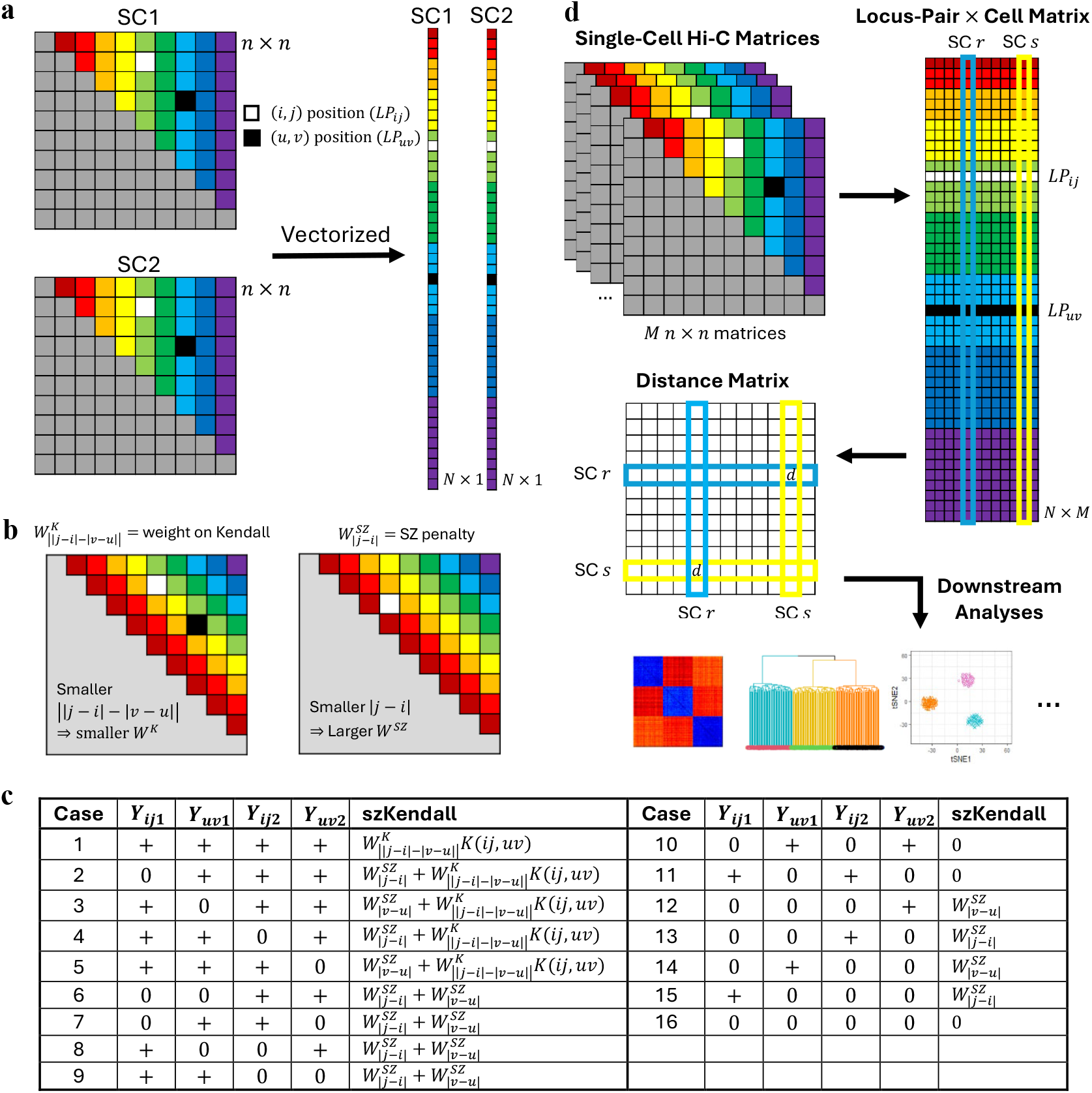
Overview of szKendall. (a) The upper triangle of an *n × n* scHi-C contact matrix is vectorized to form a vector of dimension 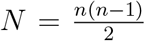. (b) Left: The band-aware weight *W*^*K*^ depends on the absolute difference between band indices of LP_*ij*_ and LP_*uv*_. Right: The structural zero penalty *W*^*SZ*^ is applied when SZ status differs across cells for a given LP. (c) SZ status combinations: Four binary SZ statuses (“0” = SZ, “+” = non-SZ) across two LPs and two cells yield 16 combinations. (d) Workflow: *M* scHi-C matrices → vectorized LPs → szKendall distance matrix (*M × M*) → downstream analysis.

However, this standard approach has two key limitations in the context of scHi-C data: (1) it ignores spatial information inherent in the 2D contact matrices; and (2) it treats all zeros equally, failing to distinguish between structural zeros and dropouts. To address the first issue, szKendall introduces band-aware weighting. Each LP resides on a “band” along the diagonal direction, defined by the genomic distance between loci (i.e., |*j* −*i*| for LP_*ij*_). For a pair of LPs, we assign a weight to their standard Kendall’s tau contribution based on the absolute difference between their band indices. Specifically, smaller differences receive lower weights (Fig. 1b, left panel), under the assumption that LPs with similar genomic distance have comparable contact counts and thus discordance may happen by chance.

To resolve the second issue, we introduce a structural zero discrepancy penalty. If a given LP is a structural zero in one cell but not in the other, this discrepancy contributes an additional penalty. Importantly, this penalty is modulated by genomic distance: discrepancies at shorter distances (i.e., bands closer to the main diagonal of the 2D contact matrix) are more informative and thus penalized more heavily (Fig. 1b, right panel). This reflects the biological intuition that contacts are more likely at short distances, so missing such contacts is more consequential. Together, these two enhancements yield the szKendall distance, a measure that leverages both spatial structure and SZ status. Note that this metric is not affected by differences in sequencing depths; thus, data across single cells do not need to be sequencing-depth normalized.

There are 16 possible combinations of SZ statuses when comparing two LPs (LP_*ij*_ and LP_*uv*_) across two single cells (SC1 and SC2), as illustrated in Fig. 1c. There, “0” denotes a structural zero, and “+” represents either a true contact or a dropout. Using szKendall, we compute a full *M* × *M* distance matrix for *M* single cells, which can then be used for visualization (e.g., heatmaps) and downstream analyses such as clustering (Fig. 1d).

In addition to the main szKendall formulation, we explore two variants: szKendall1, which assigns greater weight to LP pairs with smaller band differences (opposite of szKendall); and szKendall2, which applies a constant weight of 1 to all LP pairs, effectively removing band information from the weighting scheme. Complete details of all three versions of szKendall are provided in the Methods Section 4.1.

### 2.2 szKendall separates subtypes by capturing salient features in scHi-C data

We evaluated the performance of szKendall and its two variants (szKendall1 and szKendall2) on three simulated datasets, Sim1, Sim2, and Sim3 (Methods Section 4.2). Each of the datasets is composed of 150 single-cell contact matrices evenly divided among three cell subtypes: ST1, ST2, and ST3. The datasets differ in the abundance and pattern of SZs, both at the subtype and individual cell levels. Specifically, Sim1 contains the highest proportion of subtype-specific SZs (that is, these SZs are common among all cells within the subtype but not across subtypes), especially for ST3. Sim2 has a lower sequencing depth and fewer subtype-specific common SZs but a larger proportion of cell-specific SZs. Sim3 exhibits the lowest proportion of both subtype- and cell-specific SZs, making it the most challenging for subtype separation.

We applied szKendall and its two variants using known SZ annotations. For comparison, we also evaluated Euclidean and Kendall distances, applied to the original contact matrices and to data preprocessed by GK and RW. The resulting pairwise distance matrices and two-dimensional t-SNE projections are shown in Fig. 2.

**Fig. 2:**
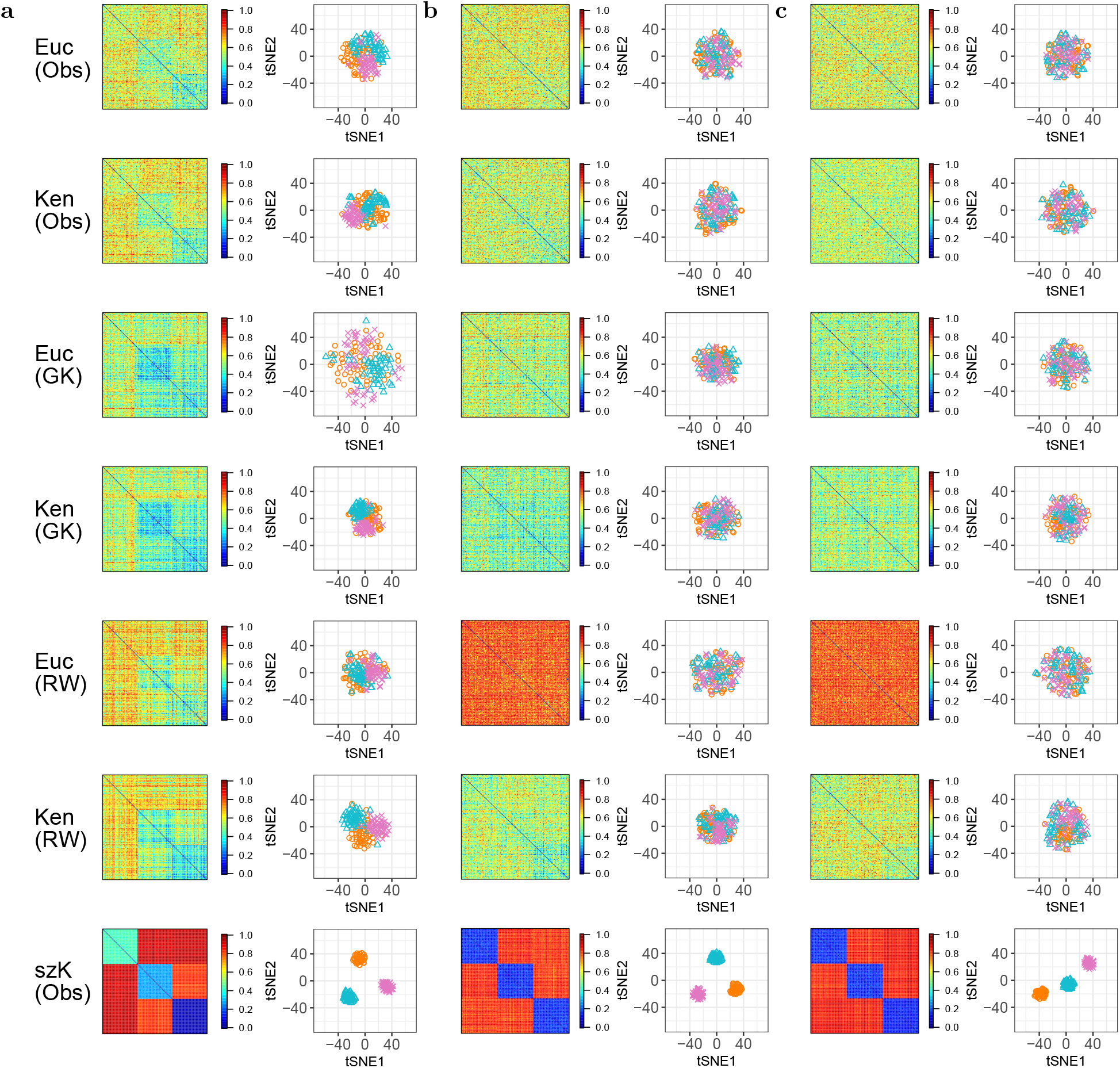
Heatmap of pair-wise cell distance matrices and t-SNE projection. The results for Sim1, Sim2, and Sim3 are provided in (a) columns 1 − 2, (b) columns 3 − 4, and (c) columns 5 − 6, respectively. Results from szKendall1 and szKendall2 are qualitatively the same as those from szKendall and are omitted for brevity. Euc = Euclidean; Ken = Kendall; szK = szKendall. Obs = “observed” counts; GK = Gaussian-kernel smoothed data; RW = random-walk smoothed data.

In Sim1, both Euclidean and Kendall distances provided discernible, albeit vague, information for separating the three subtypes. However, their performance degraded when data smoothing (GK or RW) was applied; these methods struggled particularly with detecting ST1. In contrast, all three szKendall methods — by incorporating SZ information and spatial structures — produced clear separation of subtypes, both in the distance heatmaps and the t-SNE projections.

The task became more challenging in Sim2, where the reduction in common SZs among cells of the same subtype weakened the signal used to distinguish cell groups. This trend continued in Sim3, where common SZs accounted for only 1.8% of all LPs (Table 1). As expected, this led to weaker subtype separation for all methods, though szKendall and its variants still outperformed the others by leveraging SZ patterns even though they are sparse.

**Table 1:**
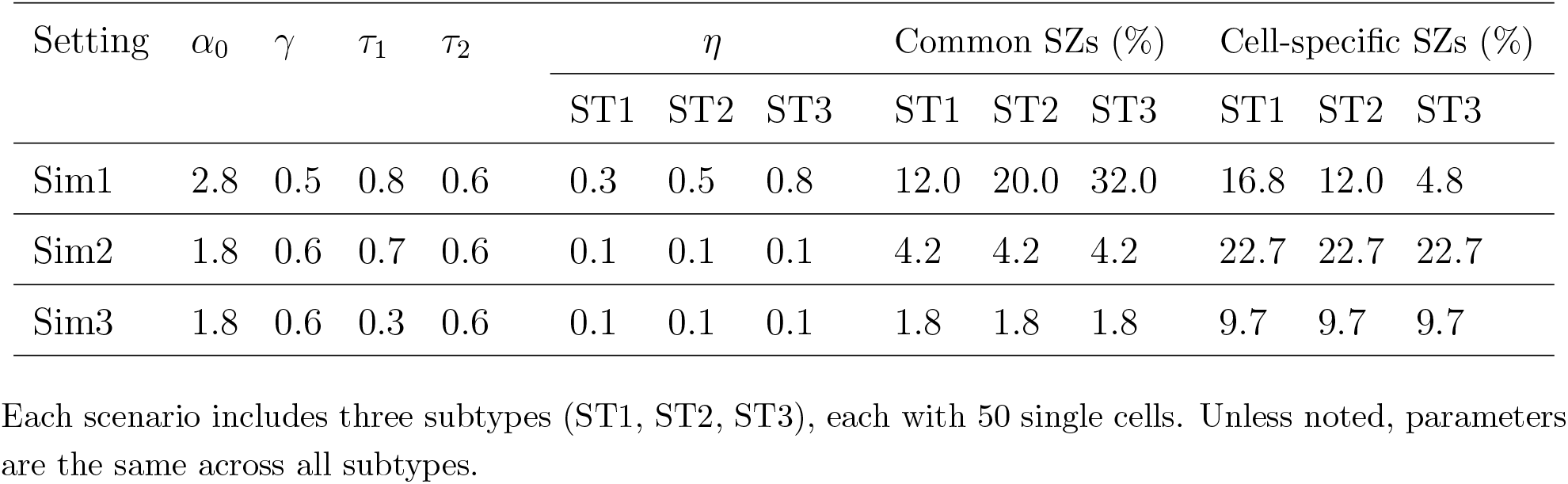
Parameter settings for three simulation scenarios (Sim1, Sim2, Sim3).

| Setting | $\alpha_0$ | $\gamma$ | $\tau_1$ | $\tau_2$ | $\eta$ | | | Common SZs (%) | | | Cell-specific SZs (%) | | |
| --- | --- | --- | --- | --- | --- | --- | --- | --- | --- | --- | --- | --- | --- |
|  |  |  |  |  | ST1 | ST2 | ST3 | ST1 | ST2 | ST3 | ST1 | ST2 | ST3 |
| Sim1 | 2.8 | 0.5 | 0.8 | 0.6 | 0.3 | 0.5 | 0.8 | 12.0 | 20.0 | 32.0 | 16.8 | 12.0 | 4.8 |
| Sim2 | 1.8 | 0.6 | 0.7 | 0.6 | 0.1 | 0.1 | 0.1 | 4.2 | 4.2 | 4.2 | 22.7 | 22.7 | 22.7 |
| Sim3 | 1.8 | 0.6 | 0.3 | 0.6 | 0.1 | 0.1 | 0.1 | 1.8 | 1.8 | 1.8 | 9.7 | 9.7 | 9.7 |

Each scenario includes three subtypes (ST1, ST2, ST3), each with 50 single cells. Unless noted, parameters are the same across all subtypes.

These results highlight that common SZs are key features for subtype discrimination. As their prevalence decreases from Sim1 to Sim2 to Sim3, subtype separation becomes progressively more difficult. Nevertheless, the szKendall framework remains effective, demonstrating robustness even under limited shared structural signal. In addition to t-SNE, we also evaluated UMAP for 2D visualization; results were qualitatively similar and are omitted here for brevity.

### 2.3 szKendall leads to better clustering criterion values

We evaluated the clustering performance of the szKendalls and two standard distance metrics, Euclidean and Kendall, using three common clustering methods: K-means (applicable only with Euclidean distance), Partitioning Around Medoids (PAM), and hierarchical clustering (the latter two are applicable to any dissimilarity measure). This results in 11 “clustering algorithms”, that is, 11 clustering method–distance measure combinations. Clustering performance was assessed using four criteria (Methods Section 4.3): adjusted Rand index (ARI), the proportion of within-cluster sum of squares to total sum of squares (WSS%), average silhouette width (ASW), and minimum isolation score (MIS).

In line with the visualization results presented in Section 2.2, ARI scores were highest for Sim1 across all methods (Fig. 3, top-left panel). Interestingly, K-means achieved near-perfect clustering for Sim1, comparable to szKendall-based methods. However, its performance substantially declined for Sim2 and Sim3, where subtype-specific SZ patterns were weaker and less prevalent. In contrast, the szKendall-based distances consistently yielded superior clustering performance across all datasets, particularly in the more challenging settings of Sim2 and Sim3.

**Fig. 3:**
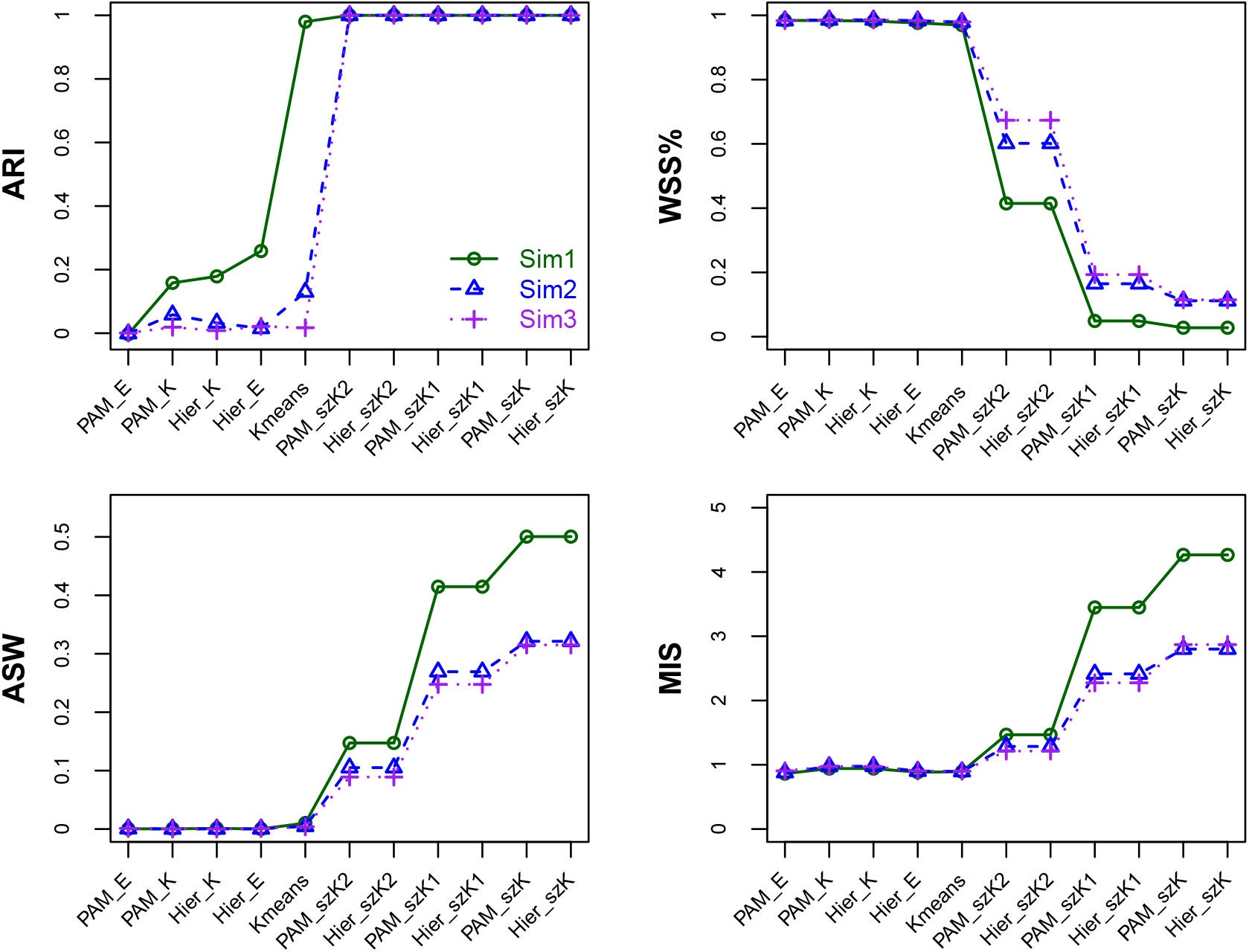
Comparison of performance of 11 algorithms of clustering-distance combinations based on four criteria. Results for all three datasets are shown, with the algorithms ordered according to increasing ARI value for Sim1. The four panels (from left to right, top to bottom) corresponds to adjusted rank index, proportion of within clusters sum of squares relative to the total sum of squares, average silhouette width, and minimum isolation score, respectively. The distance measures considered are: Euclidean (E), Kendall (K), and szKendall(1)(2) (szK, szK1, szK2). The clustering algorithm considered are K-means (with Hartigan-Wong algorithm), PAM, and hierarchical (Hier).

Despite K-means performing well on Sim1 in terms of ARI, it failed to produce well-separated clusters as indicated by the WSS% values (Fig. 3, top-right panel). The within-cluster sum of squares was nearly equal to the total sum of squares, suggesting limited separation. Moreover, it showed little advantage over other non-SZ-aware algorithms under this evaluation criterion. In contrast, all three szKendall variants, when combined with PAM or hierarchical clustering, resulted in substantially lower WSS% values across all three datasets, particularly for szKendall and szKendall1.

With regard to ASW, a higher value indicates better intra-cluster cohesion and inter-cluster separation. Again, szKendall-based approaches outperformed traditional methods, further confirming their effectiveness in identifying well-separated cell subtypes (Fig. 3, bottom-left panel). A similar trend was observed with MIS, which assesses cluster separation from a different perspective (Fig. 3, bottom-right panel). None of the non-SZ-aware algorithms attained an isolation score above 1, indicating inadequate separation (Supplementary Table S1). In contrast, szKendall and szKendall1 consistently produced MIS values that are much larger than 1, reflecting tight clusters that are well separated from one another.

It is worth noting that szKendall2, which does not incorporate spatial information in its weighting of discordant LP pairs, like Euclidean and standard Kendall, consistently lagged behind szKendall and szKendall1. This underscores the importance of incorporating spatial context in accurately quantifying dissimilarities between contact matrices.

To assess the robustness of clustering performance under challenging conditions, we focused on Sim3, the most difficult setting characterized by the lowest proportions of SZs, both common within subtypes and cell-specific. We evaluated the variability of clustering outcomes by repeating the simulation procedure 100 times. In each iteration, we computed clustering performance using both the original dissimilarity matrices and their corresponding 2D projections obtained via t-SNE or UMAP.

The results demonstrate that all six clustering algorithms based on szKendall variants exhibited highly consistent performance across runs, with minimal variability in the evaluation criteria (Supplementary Fig. S2). For the remaining five algorithms that do not utilize the SZ information nor the spatial features in the distance calculation, the results are consistently worse than the szKendall-based algorithms, with the degrees of degradation similar to those seen in Fig. 3. This shows that the much superior performance of szKendall and szKendall1 presented in Fig. 3 was not due to a particularly favorable dataset. Further, dimensionality reduction did not substantially alter clustering performance for all methods, as it is clearly seen that the szKendall-based approaches consistently outperformed the other algorithms across all datasets.

### 2.4 szKendall performs well under a wide range of SZ misspecifications

As demonstrated earlier, the szKendall methods produced near-perfect clustering results with excellent subtype separation, even under the most challenging scenario in Sim3, where the proportion of SZs is low. These results, however, were based on the idealized assumption that all SZs were correctly specified. In practical applications, SZs must be inferred using algorithms such as HiCImpute [21] and scHiCSRS [22], which inevitably exhibit varying levels of sensitivity (correctly identifying true SZs) and specificity (correctly identifying true DOs).

To assess the robustness of the szKendall dissimilarity measures under imperfect SZ annotations, we conducted a study using Sim3 data under 42 combinations of sensitivity and specificity. In the first 21 settings, we fixed the sensitivity level (*α* = 0.95, 0.8, or 0.6) by randomly selecting an *α* proportion of true SZs to retain as correctly identified. Then, we varied the specificity (*β*) from 0.9 to 0.3 in decrements of 0.1 by randomly selecting a *β* proportion of DOs to retain as correctly labeled. For each configuration, the resulting data were analyzed using the szKendall distance metrics with PAM clustering, and the process was repeated 100 times to evaluate variability. Results using hierarchical clustering were qualitatively similar and are omitted for brevity. In the other 21 settings, we fixed the specificity and varied the sensitivity in a similar manner.

When both sensitivity and specificity were at least 80%, all three szKendall dissimilarity measures continued to reach near-perfect clustering performance. This was reflected in high ARI, low WSS%, high ASW, and MIS exceeding 1 (Fig. 4 and Supplementary Fig. S3). As sensitivity or specificity decreased, clustering performance began to decline, particularly for szKendall2. Nonetheless, even with only 60% sensitivity and 40% specificity (or vice versa), the szKendall methods consistently outperformed PAM clustering based on Euclidean or Kendall distances across all four evaluation criteria.

**Fig. 4:**
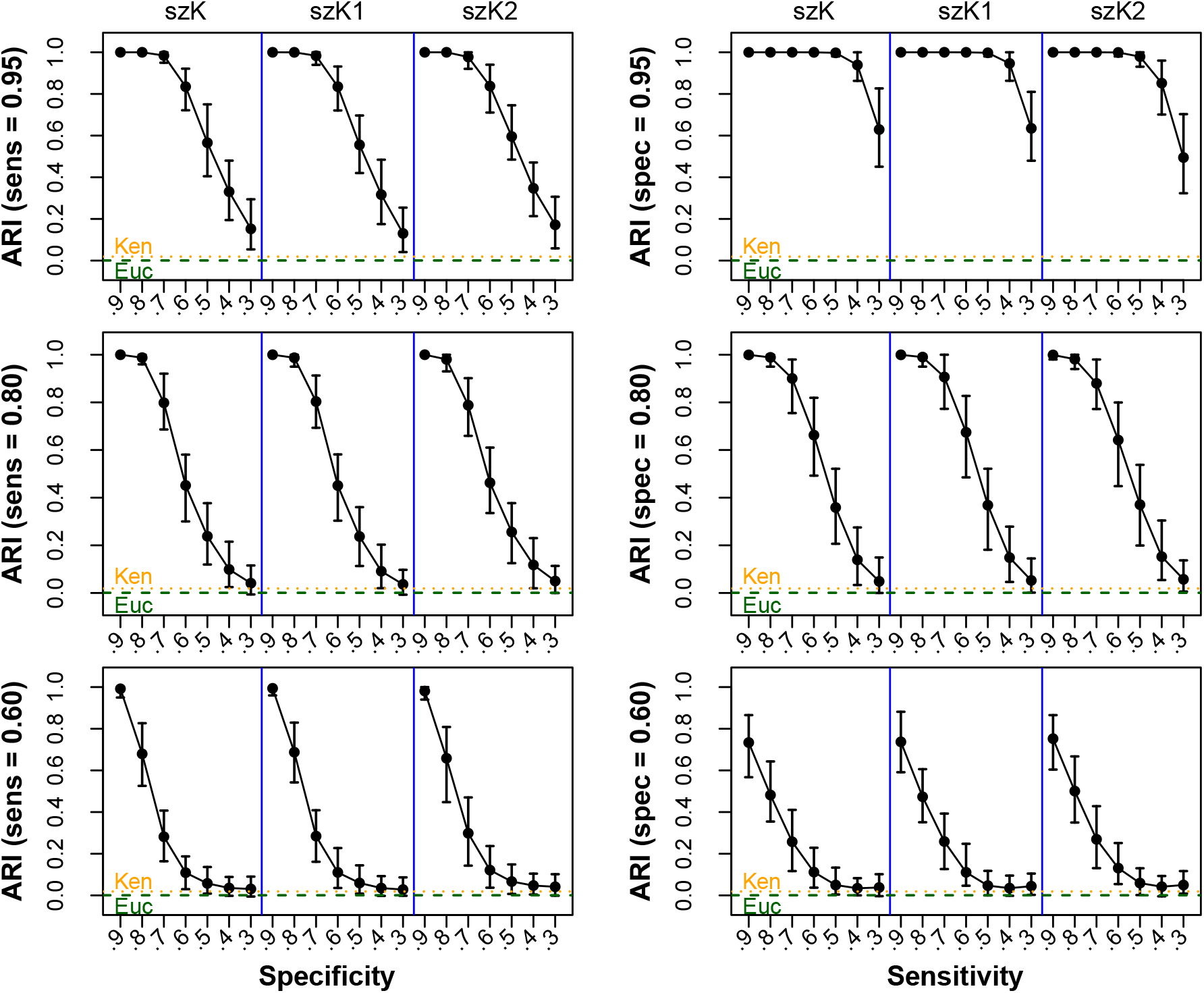
Performance evaluation of the three szKendall distance measures under reduced sensitivity and specificity. For each specified sensitivity and specificity setting, the bar provides a 95% confidence interval of the ARI value, with the curve connecting the means. The horizontal dashed lines indicate the ARI values for Kendall (orange: 0.0189) and Euclidean (green: 0.0006).

### 2.5 szKendall better delineates cell phases in mESC

We evaluated the performance of the proposed distance measures using a mouse embryonic stem cell (mESC) scHi-C dataset at 1 Mb resolution. The dataset, processed by scHi-CSim [25] (https://github.com/zhanglabtools/scHi-CSim/tree/main/data), includes 21 real single cells (GEO accession: GSE94489), each paired with a simulated cell, resulting in a total of 42 cells. These cells span three distinct stages of the cell cycle: 30 in G1 phase, 4 in early-S phase, and 8 in late-S/G2 phase. Unlike the balanced group sizes in our earlier simulation studies, this real dataset is notably imbalanced.

To prepare the data, we first scaled the sequencing depth such that each cell had approximately 1 million read pairs, reflecting a typical scHi-C experiment [12, 26]. To better mirror biological patterns, we distributed 90% of the reads to intra-chromosomal contacts [27, 28]. Zero-count LPs were then annotated as structural zeros (SZs) or dropouts (DOs) (Methods Section 4.4), resulting in approximately 7% of LPs being labeled as SZs.

We applied five distance measures — Euclidean, Kendall, szKendall, szKendall1, and szKendall2 — to compute pairwise dissimilarities among the 42 cells. Visual inspection of the resulting distance matrices and their UMAP projections (Fig. 5a) reveals that cells are poorly separated when using Euclidean or Kendall’s tau distances. In contrast, szKendall and szKendall1 attain much clearer separation of the three cell-cycle stages, while szKendall2 performs somewhat less effectively.

**Fig. 5:**
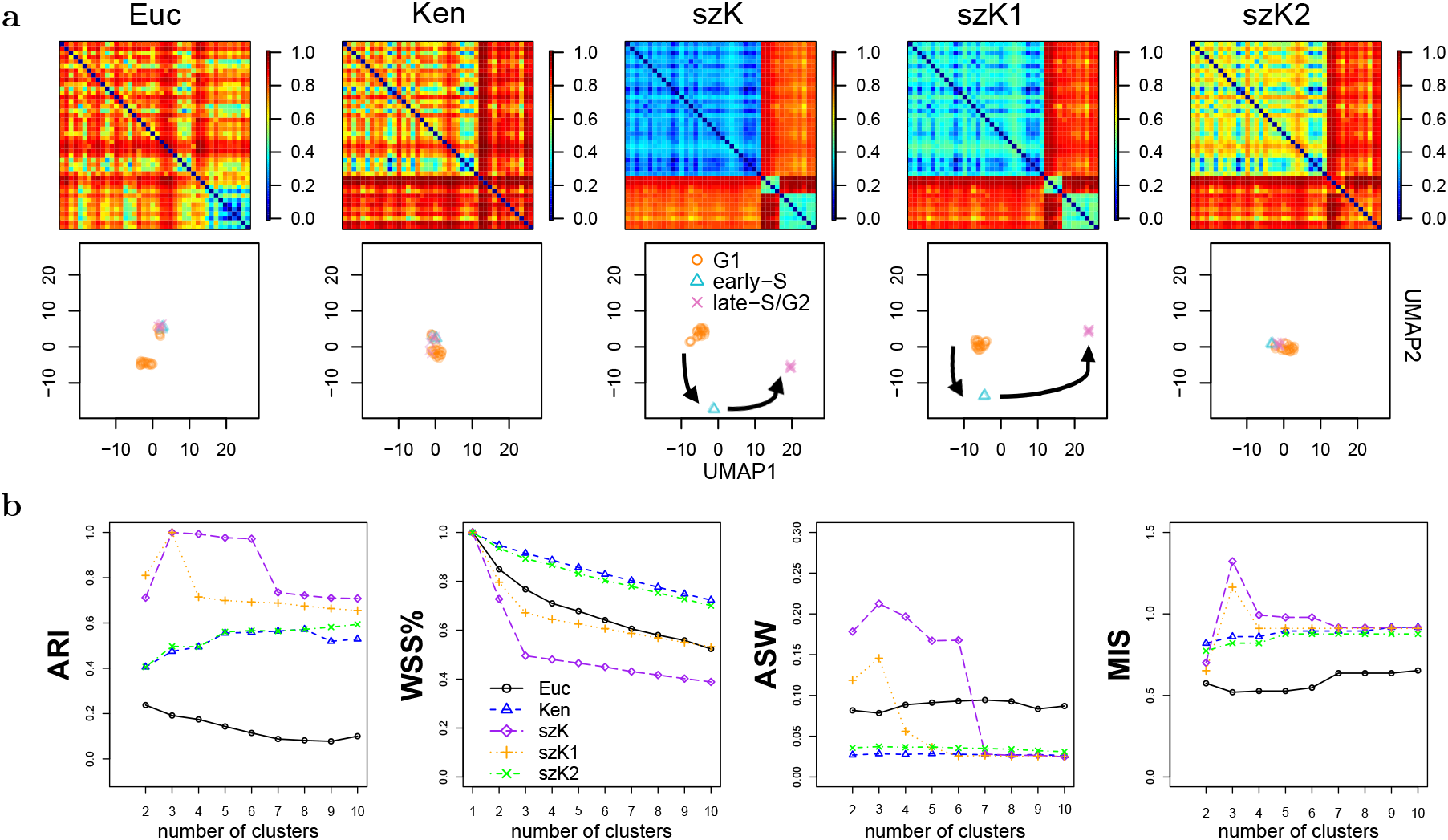
Results from analyzing the mESC data using five dissimilarity measures. (a) Heatmap of distance matrices and UMAP projections; (b) Clustering assessment criterion values for PAM clustering over the range of 2–10 clusters.

These findings were further supported by clustering analysis using the PAM algorithm applied to each of the five dissimilarity matrices. All four clustering evaluation criteria consistently confirmed the superiority of SZ-aware distances, especially szKendall and szKendall1 (Fig. 5b). Notably, for both of these methods, WSS% and ASW indicate the optimal number of clusters to be *k* = 3, and at *k* = 3, the ARI reaches 1, signifying perfect clustering of cells into their respective cell-cycle stages. Results based on hierarchical clustering agreed with the ones based on PAM (Supplementary Fig. S4).

### 2.6 szKendall separates two neuronal subtypes in prefrontal cortex cells

We analyzed a sn-m3C-seq dataset containing scHi-C profiles from human prefrontal cortex cells (https://github.com/dixonlab/scm3C-seq), focusing on two excitatory neuronal subtypes, L4 and L5, which reside in distinct cortical layers. Previous studies have shown that cells from these subtypes do not separate well using the raw scHi-C data [23, 21]. However, using the imputed data from HiCImpute, which identifies SZs and imputes DO values, led to a clear separation of L4 and L5. Most interestingly, two further subtypes within each of L4 and L5 were also revealed using the K-means clustering method on Euclidean distances computed from the improved data [21].

Motivated by these findings, we investigated whether the proposed szKendall distances, when combined with HiCImpute, could also reveal the separation of L4 and L5 subtypes in this challenging dataset. We also assessed whether the additional substructure observed within L4 and L5 could simply be an artifact of the Euclidean distance or could also be replicated by the SZ-aware szKendalls. Our expectation of good performance from szKendalls was informed by the strong sensitivity and specificity acquired by HiCImpute [21] and our simulation results (Supplementary Table S2).

Using a 5% False Positive Count Rate (FPCR) threshold (Methods Section 4.5), we classified observed zeros as SZs or DOs based on the posterior probabilities obtained from HiCImpute. Zeros labeled as SZs remained unchanged; zeros labeled as DOs were replaced by their imputed values; and positive counts were unchanged for all distance calculations. We computed five distances: Euclidean and Kendall (both on observed and imputed data), and the three szKendall variants (on imputed data with SZ annotations).

Substantial improvements in separating L4 and L5 cells after HiCImpute are evident in both the heatmaps and their projections using t-SNE and UMAP (Supplementary Fig. S5a). Notably, Euclidean distances on the imputed data revealed not only clear L4–L5 separation but also finer subdivisions into two subtypes per group, consistent with prior findings [21]. This subtype structure was not apparent with Kendall or any of the szKendall dissimilarity measures.

To further validate these observations, we applied K-means and PAM clustering methods to all seven distance matrices: Euclidean and Kendall based on the observed data, and Euclidean, Kendall, and the three szKendalls based on the improved data. Results were consistent with visualizations. In particular, all three szKendalls achieved perfect separation between L4 and L5 when *k* = 2, as shown in the confusion matrices (Table 2). Moreover, while Euclidean distances on the imputed data presents some weak evidence of a four-cluster structure based on WSS%, the remaining methods indicated little support for more than two clusters (Supplementary Fig. S5b and S6). When clustering was forced into four groups, the szKendalls yielded two dominant clusters, with the other two nearly empty (Table 2), suggesting that the two-cluster result yielded tighter and more coherent clusters than with a larger number of clusters.

**Table 2:**
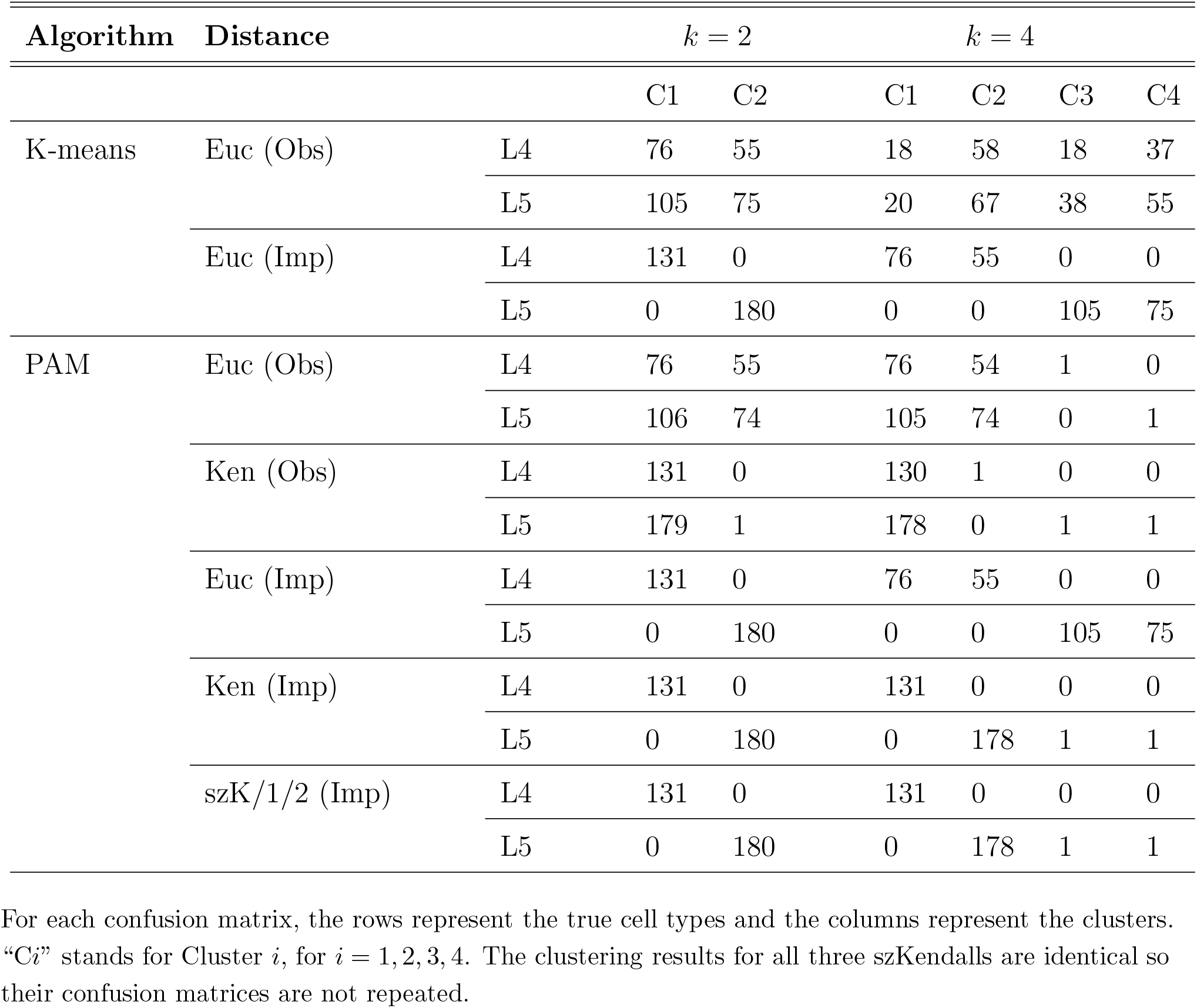
Confusion matrices of the clustering results on the prefrontal cortex data when the number of clusters is specified to be *k* = 2 or *k* = 4.

## 3 Discussion

In this study, we introduced szKendall, a novel spatial-structural-zero-aware dissimilarity measure tailored for scHi-C data. Unlike existing measures, szKendall is explicitly designed to incorporate SZ information along with DO imputed values, leveraging spatial structure in chromatin contact maps to better quantify cell-to-cell dissimilarity. The proposed measure comprises two main components: the first re-weights the classical Kendall’s tau distance to reflect spatial proximity between bands of LP pairs, while the second directly integrates inferred SZs and the proximity of the genomic locations of the two loci in each LP. To the best of our knowledge, this is the first method that explicitly incorporates SZ information into a dissimilarity measure for scHi-C analysis. We believe this often-overlooked information can significantly improve cell clustering and subtype identification.

Through extensive simulation studies, we compared szKendall with commonly used dissimilarity measures, including Euclidean and standard Kendall. When the true SZ positions were known, szKendall consistently outperformed both baselines, achieving more accurate clustering even in cases where SZs were far outnumbered by DOs (e.g., 11.5% vs. 81% in Sim3). These improvements remained visible after dimensionality reduction via t-SNE and UMAP. Note that both projections were always employed, but we may only show results of one of them for brevity, as they are qualitatively the same. Since the true SZs are typically unknown in practice, we evaluated szKendall under various levels of SZ and DO misclassification. The method remained highly robust: even with only 60% sensitivity and 40% specificity (or vise versa) in SZ detection, szKendall still surpassed Euclidean and Kendall in clustering performance.

In real data analysis, we revisited the prefrontal cortex dataset, which includes two excitatory neuronal subtypes (L4 and L5) that are challenging to separate based on raw scHi-C data from [23]. Although HiCImpute [21] identified possible substructures (two subtypes each within L4 and L5) using Euclidean distance on the imputed data, we found little support for such finer division when using szKendalls. Instead, all three variants of szKendall provided a much clearer separation between the two main subtypes, suggesting that structural zeros and their spatial information play a crucial role in recovering biologically meaningful groupings that Euclidean and standard Kendall’s tau distances may obscure.

While our results demonstrate the promise of szKendall, several limitations remain. First, the effectiveness of szKendall depends heavily on accurate SZ detection. In this study, we employed HiCImpute, which yielded strong results, but alternative SZ/DO classification algorithms may potentially lead to better performance. Second, our analyses showed that szKendall and szKendall1 performed similarly (and both outperformed szKendall2) despite using opposite weighting schemes. In szKendall, a greater weight is given to LPs with larger band differences, emphasizing more detrimental effects of short- versus long-range contact discrepancies, whereas szKendall1 places more weight on the opposite type of discrepancies. The underperformance of szKendall2, which applies a constant weight, suggests that both proximity and distance contribute important, but distinct, biological signals. A hybrid weighting scheme that balances both short- and long-range differences may yield even better clustering accuracy and warrants future exploration.

Finally, a practical limitation is the computational cost of szKendall, especially when applied to a large datasets (Supplementary document Section 1). Future work will focus on developing optimized algorithms or scalable approximations to improve runtime and memory efficiency, which will be critical for enabling broader adoption of the method in large-scale single-cell genomic studies.

## 4 Materials and Methods

### 4.1 szKendall Dissimilarity Measures

For a pair of single cells, SC1 and SC2, the standard Kendall’s tau dissimilarity quantifies the number of pairwise “discordances” in their contact counts. Let *Y*_*ijs*_ and *Y*_*uvs*_ denote the contact counts at two LPs, LP_*ij*_ and LP_*uv*_, for cell *s* = 1, 2. The standard Kendall dissimilarity for this pair of LPs is defined as:

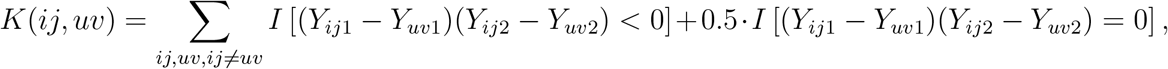

where *I*(·) is the usual indicator function. A value of 1 indicates a discordant pair, 0 a concordant one, and 0.5 a tie. We extend this framework by incorporating two additional types of information: spatial structure of chromatin contacts and inferred structural zeros (SZs). The proposed szKendall measure consists of the following two key components.

#### Weighted Kendall’s Discordance Based on band difference

When none or only one of the four contact counts (*Y*_*ij*1_, *Y*_*uv*1_, *Y*_*ij*2_, *Y*_*uv*2_) is identified as an SZ (i.e., Cases 1–5 in Fig. 1c), the standard Kendall discordance is computed and weighted according to the bandwise distance between LP_*ij*_ and LP_*uv*_. Specifically, the weight is a function of the absolute difference in the two LP bands: ||*j* −*i*| − |*v* −*u*||.

For szKendall, the weight is defined as:

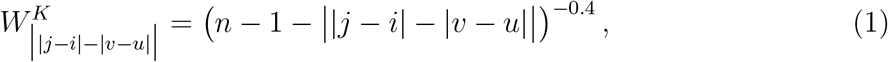

where *n* is the size of the contact matrix. This function assigns lower weights to LPs with similar genomic distances (i.e., closer bands), under the assumption that such discordances may arise due to random variation.

We also considered two variants modifying this weighting scheme. szKendall1 emphasizes discrepancies for LP pairs with similar genomic distances:

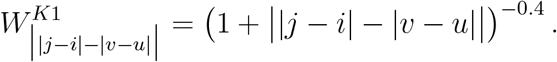

On the other hand, szKendall2 applies a constant weight:

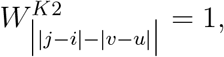

effectively reducing the weighted version to an unweighted Kendall’s tau.

#### Structural Zero Discordance Penalty

For LP pairs with at least one inconsistent SZ status (Cases 2 – 9 and 12 – 15 in Fig. 1c), we introduce an additional penalty term. If the SZ assignment differs between cells in LP_*ij*_, we add a dissimilarity penalty that scales with the proximity of the LP to the main diagonal of the contact matrix:

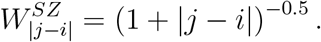

This formulation gives larger penalties to discrepancies near the diagonal, where contact counts are typically higher unless suppressed by true structural zeros. The same penalty is applied independently for discrepancies in LP_*uv*_.

The final szKendall dissimilarity between two cells is computed by aggregating the weighted discordances and SZ penalties across all LP pairs. This integrated approach allows us to better capture biologically meaningful differences while accounting for both spatial organization and dropout artifacts in scHi-C data.

### 4.2 Simulation Procedure

Each simulated dataset consists of 150 scHi-C contact matrices representing three distinct “subtypes” of 50 cells each. These subtypes are derived based on the 3D structure of a segment in chromosome 19 (61 loci) from a K562 cell (accession: GSM2109974) obtained from publicly available data [11] (https://www.ncbi.nlm.nih.gov/geo/query/acc.cgi?acc=GSE80006). Following the approach in prior studies [29, 21], this setup mimics subtype heterogeneity within a homogeneous cell population. The simulation process involves four key steps:

#### Step 1: Compute Euclidean distances

We calculate the Euclidean distance *d*_*ij*_ between every LP_*ij*_, *i* < *j* = 2, 3, · · ·, *n* using the 3D coordinates of the K562 cell. For the diagonal elements, we set *d*_*ii*_ = 0.1 for all *i* = 1, *…, n*, which is slightly smaller than the smallest nonzero distance observed among distinct loci of the K562 cell. This nonzero value facilitates modeling interaction intensities using an additive structure after log transformation.

#### Step 2: Generate expected contact matrix

We define the expected contact matrix 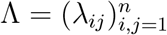, where the log-transformed expected contact is modeled as:

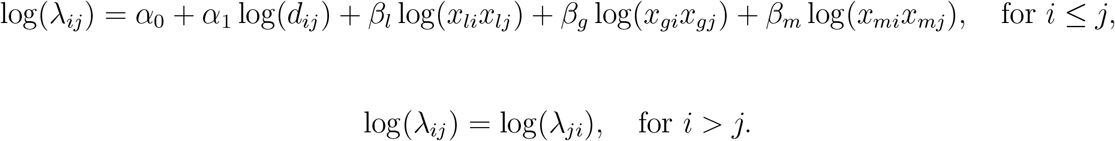

Here, *α*_0_ is a global scaling parameter controlling sequencing depth, and *α*_1_ = −1 reflects the biophysical law, stipulating that contact frequency decreases with spatial distance [30]. The covariates *x*_*li*_, *x*_*gi*_, *x*_*mi*_ mimic fragment length, GC content, and mappability, respectively, and are independently sampled with *x*_*li*_ ~ Uniform(0.2, 0.3), *x*_*gi*_ ~ Uniform(0.4, 0.5), and *x*_*mi*_ ~ Uniform(0.9, 1), and with corresponding coefficients *β*_*l*_ = *β*_*g*_ = *β*_*m*_ = 0.9.

#### Step 3: Introduce structural zeros (SZs)

To introduce both within-subtype shared and cell-specific SZs, we proceed as follows:

1. *Identify SZ candidates:* For each *λ*_*ij*_ in the upper triangular of Λ, if it falls in the bottom *γ ×* 100 percentile, it becomes an SZ candidate with probability *τ*_1_.
2. *Assign common SZs within subtypes:*

- For each subtype, a proportion *η* of the SZ candidates are randomly selected as *common* SZs shared by all 50 cells in that subtype.
- The remaining (1 −*η*) proportion of the SZ candidates are subject to cell-specific assignment: each is assigned as an SZ in a given cell with probability *τ*_2_.
- *Define cell-specific expected contact matrix:* For each cell *m*, define its expected matrix 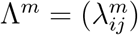, where

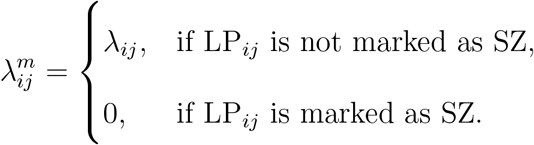

This setup results in each cell having an expected SZ proportion of *γ* ·*τ*_1_ · [*η* + (1 −*η*) ·*τ*_2_].

#### Step 4: Simulate observed contact matrices

For each cell *m*, we generate the observed count matrix 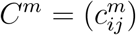 by sampling:

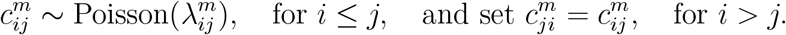

If 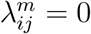, then 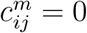 with probability 1. The resulting matrix is symmetric.

In our simulation study, we consider three scenarios (Sim1, Sim2, Sim3) by varying key parameters, as summarized in Table 1. Each dataset consists of three subtypes with 50 cells each. The table also shows the expected proportions of common and cell-specific SZs in each subtype. Compared to Sim1, Sim2 has a lower sequencing depth parameter (*α*_0_) and a smaller proportion of common SZs (*η*). However, the overall proportion of SZ candidates (*γ* ·*τ*_1_) remains similar to Sim1 (40% in Sim1 versus 42% in Sim2). Comparing to Sim2, Sim3 has an even smaller *τ*_1_, reducing the number of SZ candidates and hence fewer SZs overall, while keeping the common vs. cell-specific ratio unchanged.

### 4.3 Four Clustering Assessment Criteria

We evaluated the performance of 11 combinations of dissimilarity measures (Euclidean, Kendall, szKendall, szKendall1, and szKendall2) and clustering methods (K-means, PAM, and hierarchical clustering) using four different criteria. Note that K-means is only applicable with Euclidean distance, while PAM and hierarchical clustering can be used with any of the dissimilarity measures. Among the four evaluation criteria, ARI requires knowledge of the true group labels. In contrast, the other three criteria assess aspects of within cluster compactness and between-cluster separation, making them interpretable when the true labels are unknown, a realistic setting for many real data applications. Additionally, for the analysis of prefrontal cortex L4 and L5 subtypes, we assessed clustering performance using both the original observed data and the Hi-C impute-enhanced data.

#### Adjusted Rand Index (ARI)

This metric quantifies the similarity between true groupings and clustering results [31, 32]. Its values range from −1 to 1, where 1 indicates perfect agreement; that is, the number of clusters matches the true number of groups and the cluster assignments align exactly with the known labels. A value near 0 suggests that the clustering is no better than random guess, while a negative value indicates worse-than-random agreement, implying that the clustering is less similar to the true grouping than would be expected by chance.

#### Within-cluster sum of squares percentage (WSS%)

This metric quantifies the proportion of the total sum of squares explained by the within-cluster sum of squares [33, 34]. The total sum of squares is calculated as the sum of all pairwise squared Euclidean distances among cells in the dataset, normalized by twice the total number of cells. For each cluster, the within-cluster sum of squares is computed similarly, using pairwise squared Euclidean distances among cells within the cluster, normalized by twice the number of cells in that cluster. Summing these values across all clusters gives the overall within-cluster sum of squares. When extending this definition to alternative measures — such as Kendall’s tau or szKendall — the squared Euclidean distance is substituted with the corresponding dissimilarity metric. The final value is expressed as a percentage between 0% and 100%, where lower values indicate greater cluster compactness or tightness.

#### Average silhouette width (ASW)

The silhouette value of an observation measures how well it fits within its assigned cluster (cohesion) compared to its potential fits in other clusters (separation) [35]. This value ranges from −1 to 1, where a higher value indicates that the observation is well matched to its own cluster and would be poorly matched to neighboring clusters. The ASW is the mean of the silhouette scores across all observations. A higher ASW suggests better within-cluster cohesion and greater separation between clusters, indicating more well-defined clustering.

#### Minimum Isolation Score (MIS)

For each cluster produced by a clustering algorithm, the isolation score is defined as the ratio of separation to diameter [36]. Separation refers to the minimum dissimilarity between any cell in the cluster and any cell in other clusters, while diameter is the maximum pairwise dissimilarity among cells within the cluster. A cluster is considered isolated if its separation exceeds its diameter; that is, if the isolation score is greater than 1. To assess overall cluster separation, we use the minimum isolation score across all clusters. Note that if a cluster contains only a single observation, its isolation score is defined as infinity.

### 4.4 Preparation of mESC data

For a mouse embryonic stem cell (mESC) scHi-C experiment at 1 Mb resolution, a typical sequencing depth is approximately 1 million read pairs per cell. This depth provides sufficient coverage to detect chromatin interactions across the genome, while still accounting for the intrinsic sparsity of scHi-C data [12, 26]. To simulate this, we first scaled the data such that the total sequencing depth per cell across the entire genome is approximately 1 million. Empirical studies show that at least 90% of reads are intra-chromosomal [27, 28]. Therefore, 90% of the total reads were allocated to intra-chromosomal interactions. In our analysis, we focused exclusively on chromosome 1. The corresponding scaling factor was computed as:

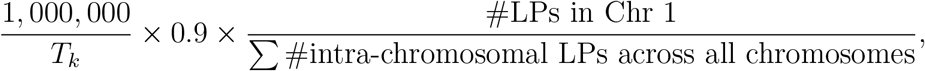

where the factor 0.9 reflected the estimated proportion of intra-chromosomal interactions and *T*_*k*_ was the observed contact counts among chromosome 1 loci for cell *k, k* = 1, 2, *…,* 42. To evaluate the performance of szKendall and its two variants, we generated ground-truth SZ positions. The procedure was as follows: for each contact count, we applied a smoothing operation using a diamond-shaped neighborhood of radius 3. Specifically, for a given position LP_*ij*_, the neighborhood was defined as {LP_*uv*_ : |*i* −*u*| + |*j* −*v*|*≤* 3}. After smoothing, we identified candidate SZ positions among those where the smoothed count was zero across all cells at the same developmental stage. From these, 50% were randomly selected as potential SZ candidates. Among the candidates, 20% were designated as common SZs shared by all cells at the same stage. From the remaining candidates, each cell independently selected 40% to serve as cell-specific SZs. This selection scheme resulted in approximately 7% of all LPs being designated as ground-truth SZs on average.

### 4.5 Control FPCR cutoff for SZ identification

HiCImpute [21] employs a Bayesian hierarchical model to estimate the posterior probability that a given LP is an SZ. By default, when the posterior probability exceeds 0.5, the LP is classified as an SZ. Otherwise, the LP is labeled either as a DO if the observed count is zero, or a positive count (PC) if the observed count is nonzero. The choice of the 0.5 cutoff arises from a simple application of Bayes’ rule, wherein a posterior probability greater than 0.5 indicates that the SZ hypothesis is more probable than not [37]. In extremely sparse data scenarios (e.g., ~90% zeros), a large fraction of LPs may naturally have posterior probabilities above 0.5, resulting in an overcalling of SZs. Moreover, since HiCImpute jointly infers SZs across cells — that is, the inferred SZ positions are common — which may lead to a non-negligible portion of positive count positions being erroneously labeled as SZs.

To address this, we introduce a data-driven thresholding approach that replaces the fixed 0.5 cutoff by controlling the False Positive Count Rate (FPCR) to be no more than a prespecified level *α* (e.g., 5%). The key idea is motivated by the fact that most artifact-induced contacts — due to random ligation, genome misassembly, or sequencing noise — are expected to be removed during preprocessing [38]. Given current sequencing depths and technological constraints in single-cell Hi-C experiments, previous studies have found a small proportion of observed zeros to be SZs [22]. Thus, making the assumption that no more than 50% of observed zeros are true SZs, we propose to estimate an empirical SZ probability threshold (*≤ α*) such that at most 50% of observed zeros are labeled SZs, keeping the FPCR below the desired level.

## Data Availability

The szKendall R package, together with the prepared R data for the real and simulated data used in this study, are available on Github: https://github.com/osu-stat-gen/szKendall.

## Acknowledgments

This research is supported in part by a grant from the National Institute of Health R01GM114142.

## Author Contributions

SL designed the study and supervised the project. YL conducted the research. SL and YL wrote the manuscript. VJ contributed to the discussions and provided feedback.

## Competing interests

The authors declare no competing interests.

## Additional information

Correspondence and requests for materials should be addressed to SL.

## Notes

### Competing Interest Statement

The authors have declared no competing interest.

