## Supplementary Document for "Novel Spatial-Structural-Zero-Aware Dissimilarity Measures for Subtype Discovery Using Single Cell Hi-C Data"

#### 1 Computational intensity of szKendall dissimilarity measures

The computation of the three versions of the szKendall distance is implemented in `Rcpp` to improve computational efficiency. For each cell, there are $N = \frac{n(n-1)}{2}$ locus pairs (LPs), where $n$ is the number of loci within a genomic segment (e.g., a chromosome) under consideration.

To compute the szKendall distance between two cells, SC1 and SC2, all LP pairs — $\text{LP}_{ij1}$, $\text{LP}_{uv1}$, $\text{LP}_{ij2}$, and $\text{LP}_{uv2}$ — are classified into 16 groups based on the combinations of SZ (structural zero) statuses at the four involved loci. Within each group, the SZ penalty term ($W^{\text{SZ}}$) in the szKendall metric can be computed in linear time with respect to $N$.

The second component of the szKendall distance involves computing the standard Kendall's tau distance and its associated weight. A naive computation of Kendall's tau has time complexity $O(N^2)$, but more efficient algorithms exist that reduce this to $O(N \log N)$ [1].

In the case of szKendall2, where all LP pairs share a constant weight of 1, the overall computational complexity is $O(N \log N)$. However, for szKendall and szKendall1, each LP pair is assigned a weight that depends on the genomic positions of the loci, which prevents the use of the optimized algorithm. As a result, the Kendall's tau component for these variants remains computationally expensive at $O(N^2)$.

Fortunately, the positional weights depend only on the loci positions and not on the observed counts or SZ statuses. Thus, all pairwise weights can be *precomputed* once in $O(N^2)$ time and reused across all cell pairs.

Assuming there are $M$ cells in the dataset, there are $\frac{M(M-1)}{2}$ unique cell pairs. When $M$ is large, this can become a computational bottleneck. However, since the distance calculation for each cell pair is independent, the workload is highly parallelizable. In our implementation, we use the R packages *doParallel* and *foreach* to distribute these computations across multiple CPU cores.

### 2 Supplementary Figures and Tables

a

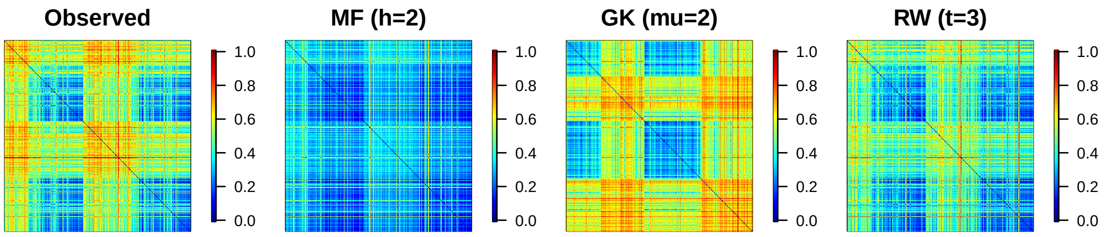


b

Observed:

| | C1 | C2 |
|---|---|---|
| L4 | 76 | 55 |
| L5 | 105 | 75 |

MF (h=2):

| | C1 | C2 |
|---|---|---|
| L4 | 76 | 55 |
| L5 | 105 | 75 |

GK (mu=2):

| | C1 | C2 |
|---|---|---|
| L4 | 63 | 68 |
| L5 | 96 | 84 |

RW (t=3):

| | C1 | C2 |
|---|---|---|
| L4 | 76 | 55 |
| L5 | 105 | 75 |

c

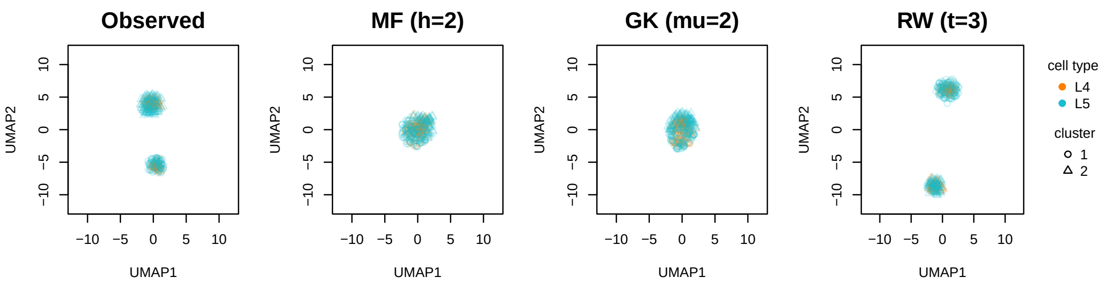


d

Observed:

| | C1 | C2 |
|---|---|---|
| L4 | 76 | 55 |
| L5 | 106 | 74 |

MF (h=2):

| | C1 | C2 |
|---|---|---|
| L4 | 77 | 54 |
| L5 | 106 | 74 |

GK (mu=2):

| | C1 | C2 |
|---|---|---|
| L4 | 77 | 54 |
| L5 | 103 | 77 |

RW (t=3):

| | C1 | C2 |
|---|---|---|
| L4 | 76 | 55 |
| L5 | 106 | 74 |

Supplementary Fig. S1: Visualization of a real data set (L4/L5) showing raw and improved data without taking SZ into account do not show clear patterns (2 main diagonal blocks in the heatmaps). (a) Heatmap visualization of the Euclidean distance matrices on the human prefrontal cortex (L4/L5) data. First heatmap is based on the raw observed data; the second heatmap is based on the mean filter (MF) smoothed counts (with smoothing parameter $h = 2$); the third heatmap is based on the Gaussian kernel (GK) smoothed data (with smoothing parameter $\mu = 2$); the last heatmap is based on the random walk (RW) with $t = 3$ steps smoothed data. (b) K-means (Hartigan-Wong algorithm) clustering result on the raw and improved data, where the number of clusters is $k = 2$. "C$i$" stands for "Cluster $i$", $i = 1, 2$. (c) UMAP 2D projection of the 311 cells, based on raw and improved data. (d) K-means (Hartigan-Wong algorithm) clustering result on projected space.

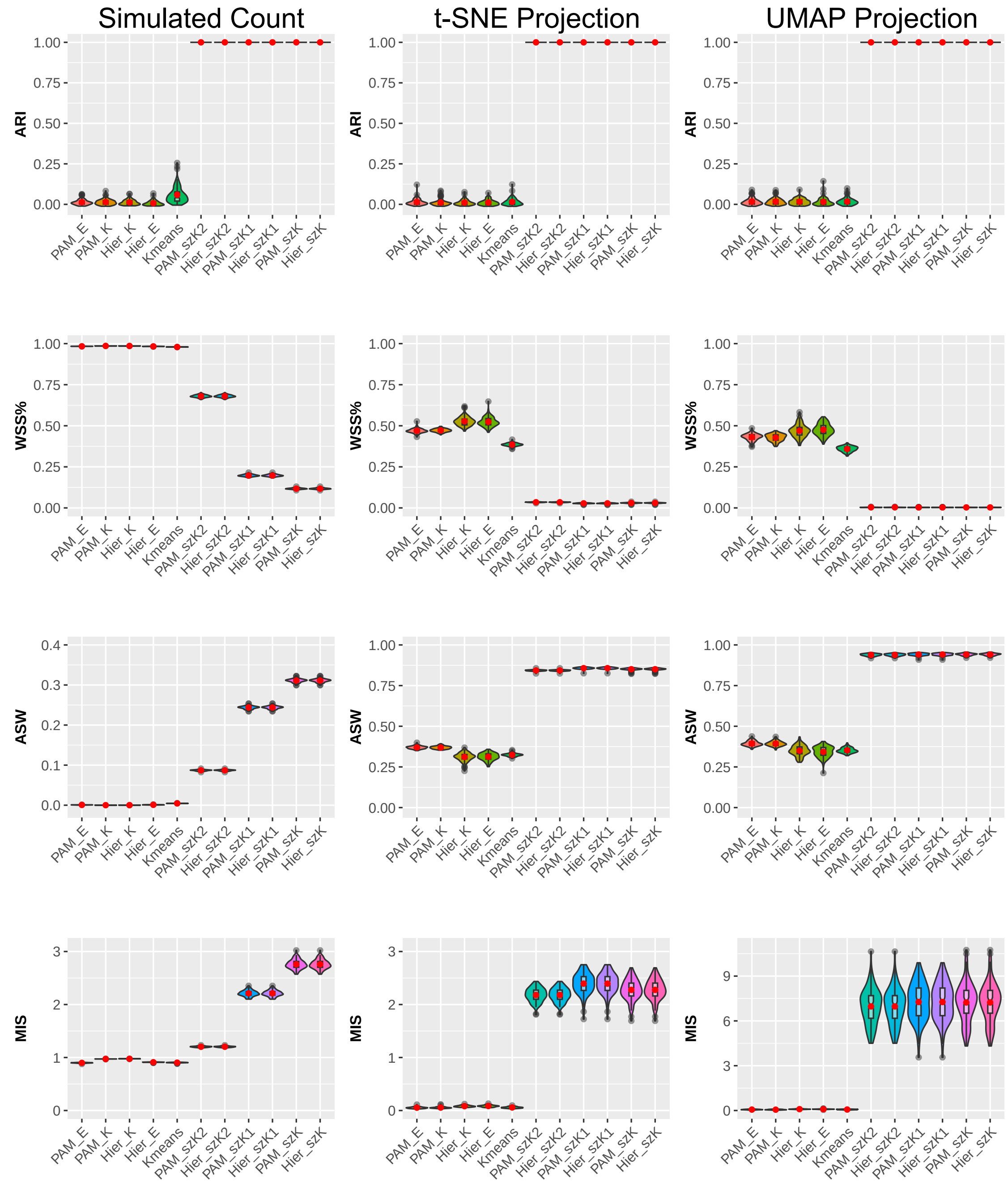


Supplementary Fig. S2: Violin plots of clustering performance over 100 Sim3 datasets evaluated by four criteria. Clustering results are based on 11 algorithms using the original 2D contact matrices (column 1), t-SNE projected data (column 2), and UMAP projected data (column 3).

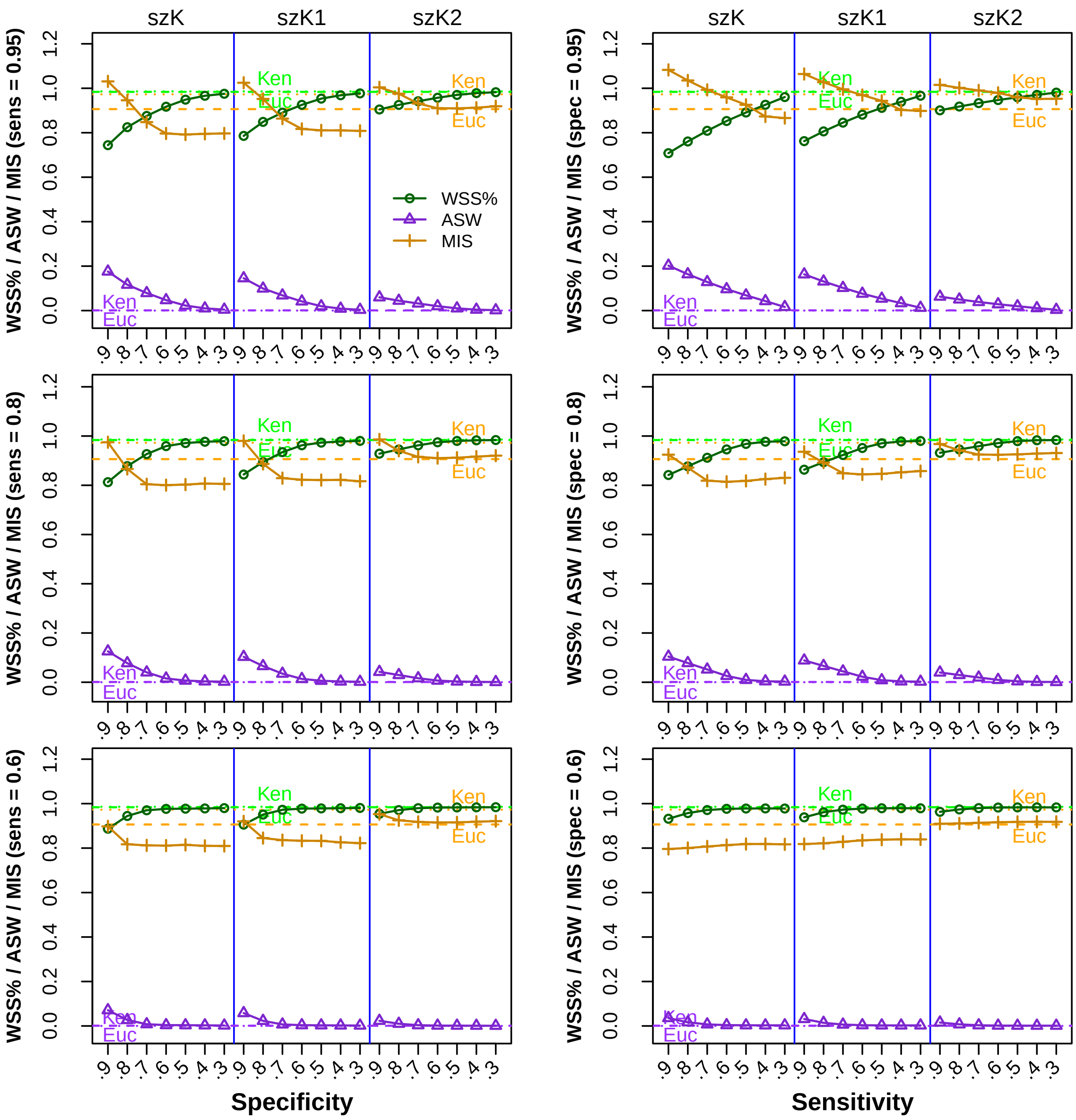


Supplementary Fig. S3: Clustering performance evaluated by criteria WSS%, ASW, and MIS in 42 settings of SZ misspecifications. Left column: Fix sensitivity and vary specificity from 0.9 to 0.3 with a decrement of 0.1. Right column: Fix specificity and vary sensitivity from 0.9 to 0.3 with a decrement of 0.1. Plotted are the mean values over 100 replications. Within each panel, the criterion values based on Euclidean and Kendall distances are marked with horizontal dashed and dotted lines, respectively.

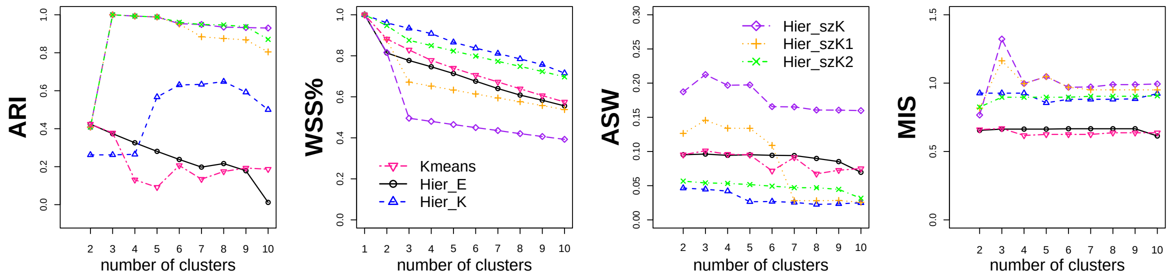


Supplementary Fig. S4: Clustering assessment criterion values for K-means and hierarchical clustering on the mESC data over the range of 2–10 clusters.

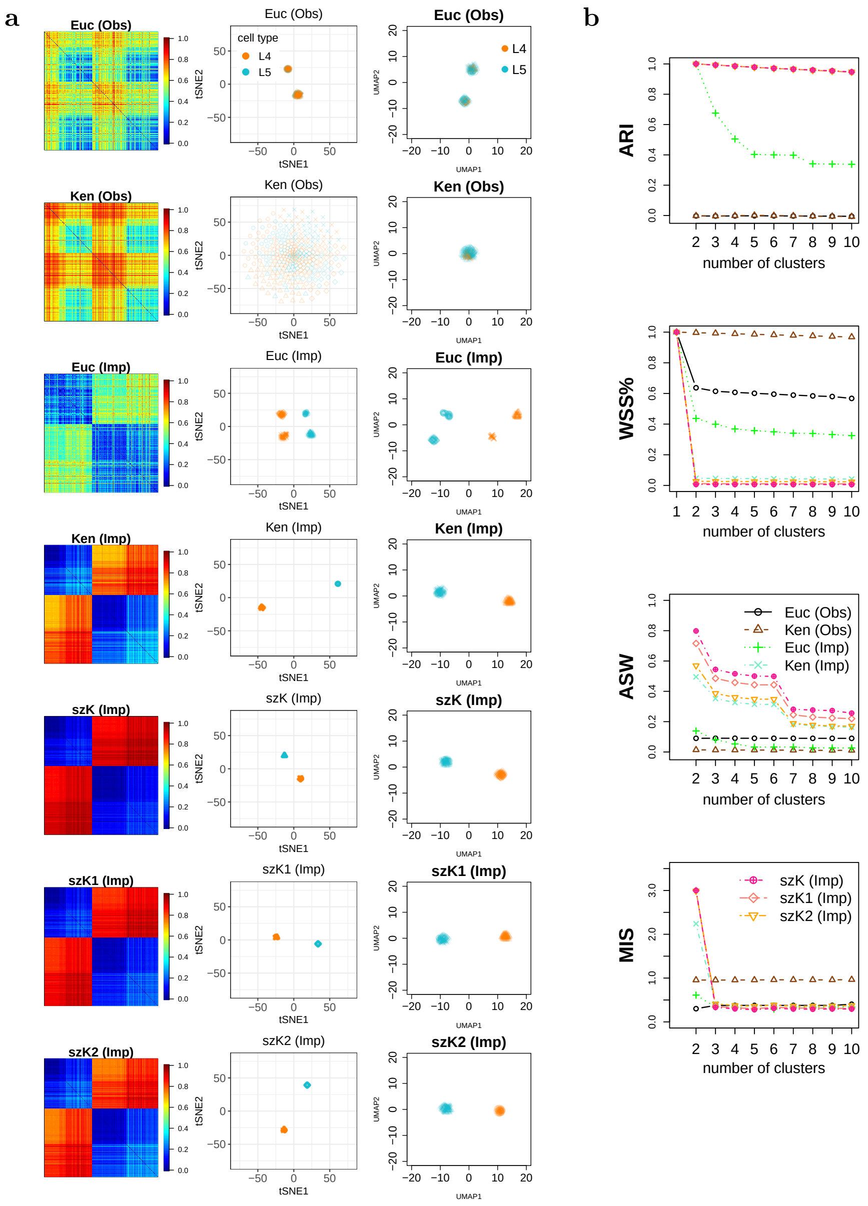


Supplementary Fig. S5: Results from analyzing the human prefrontal cortex data. (a) Column 1: Heatmap visualization of the seven distance matrices on the observed and imputed data with FPCR $\leq 5\%$ cutoff. In each distance matrix, the first 131 rows/columns represent the L4 cells, and the last 180 rows/column represent the L5 cells. Column 2: t-SNE projections. Column 3: UMAP projections. For both projections, the orange points stand for L4 cells, and light blue points L5 cells. (b) Four criterion values of PAM clustering.

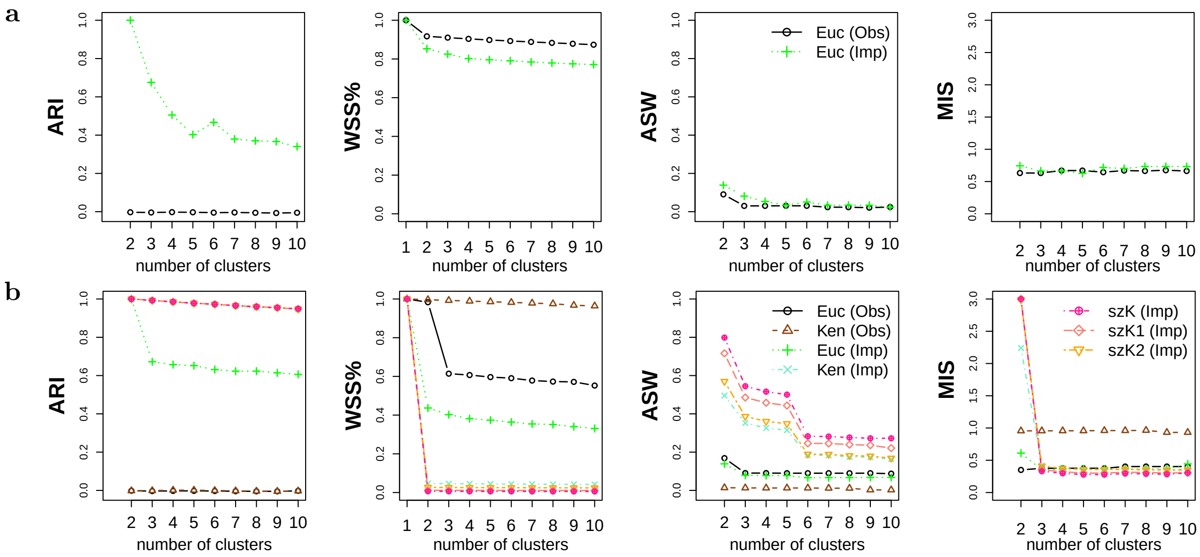


Supplementary Fig. S6: Clustering assessment criterion values of (a) K-means and (b) hierarchical clustering on the observed and imputed prefrontal cortex data over the range of 2–10 clusters.

Supplementary Table S1: Clustering performance assessment based on four evaluation criteria for three simulated datasets (Sim1, Sim2, Sim3).

| **Data** | **Distance** | **Clustering Method** | **ARI** | **WSS%** | **ASW** | **MIS** |
|---|---|---|---|---|---|---|
| Sim1 | Euclidean | K-means | 0.9799 | 0.9695 | 0.0102 | 0.8931 |
| | | PAM | -0.0034 | 0.9842 | 0.0003 | 0.8617 |
| | | Hier | 0.2585 | 0.9763 | 0.0002 | 0.8836 |
| | Kendall | PAM | 0.1583 | 0.9838 | 0.0004 | 0.9405 |
| | | Hier | 0.1787 | 0.9818 | 0.0011 | 0.9410 |
| | szKendall | PAM/Hier | 1 | 0.0279 | 0.5005 | 4.2657 |
| | szKendall1 | PAM/Hier | 1 | 0.0490 | 0.4148 | 3.4478 |
| | szKendall2 | PAM/Hier | 1 | 0.4147 | 0.1473 | 1.4672 |
| Sim2 | Euclidean | K-means | 0.1297 | 0.9788 | 0.0047 | 0.8948 |
| | | PAM | -0.0012 | 0.9835 | 0.0007 | 0.8797 |
| | | Hier | 0.0152 | 0.9834 | 0.0004 | 0.9012 |
| | Kendall | PAM | 0.0581 | 0.9859 | 0.0001 | 0.9737 |
| | | Hier | 0.0325 | 0.9859 | 0.0002 | 0.9769 |
| | szKendall | PAM/Hier | 1 | 0.1117 | 0.3213 | 2.8000 |
| | szKendall1 | PAM/Hier | 1 | 0.1647 | 0.2693 | 2.4136 |
| | szKendall2 | PAM/Hier | 1 | 0.6016 | 0.1051 | 1.2851 |
| Sim3 | Euclidean | K-means | 0.0171 | 0.9796 | 0.0039 | 0.8997 |
| | | PAM | 0.0006 | 0.9834 | 0.0010 | 0.9059 |
| | | Hierarchical | 0.0215 | 0.9837 | 0.0010 | 0.9036 |
| | Kendall | PAM | 0.0189 | 0.9859 | 0.0002 | 0.9727 |
| | | Hier | 0.0080 | 0.9857 | 0.0001 | 0.9722 |
| | szKendall | PAM/Hier | 1 | 0.1147 | 0.3151 | 2.8698 |
| | szKendall1 | PAM/Hier | 1 | 0.1932 | 0.2475 | 2.2749 |
| | szKendall2 | PAM/Hier | 1 | 0.6736 | 0.0890 | 1.2132 |

Supplementary Table S2: Sensitivity and specificity of smoothing algorithms for correctly identifying SZs and DOs. Accuracy is calculated as the proportion of correctly labeled observed zeros.

| Algorithm | Sensitivity | Specificity | Accuracy |
|---|---|---|---|
| GK | 95.6 (0.2) | 9.2 (0.4) | 19.9 (0.3) |
| RW | 88.2 (1.8) | 12.9 (2.0) | 22.3 (1.5) |
| HiCImpute | 77.7 (1.3) | 77.7 (0.4) | 77.7 (0.4) |

Each cell in the table contains the mean (SD) in percentage over 100 datasets simulated under Sim3 setting. For each smoothing/imputation algorithm, the 5% FPCR threshold is applied to infer SZs.
